# The potential for gas-free measurements of absolute oxygen metabolism during both baseline and activation states in the human brain

**DOI:** 10.1101/705186

**Authors:** Eulanca Y. Liu, Jia Guo, Aaron B. Simon, Frank Haist, David J. Dubowitz, Richard B. Buxton

## Abstract

Quantitative functional magnetic resonance imaging methods make it possible to measure cerebral oxygen metabolism (CMRO_2_) in the human brain. Current methods require the subject to breathe special gas mixtures (hypercapnia and hyperoxia). We tested a noninvasive suite of methods to measure absolute CMRO_2_ in both baseline and dynamic activation states without the use of special gases: arterial spin labeling (ASL) to measure baseline and activation cerebral blood flow (CBF), with concurrent measurement of the blood oxygenation level dependent (BOLD) signal as a dynamic change in tissue R_2_*; VSEAN to estimate baseline O_2_ extraction fraction (OEF) from a measurement of venous blood R_2_, which in combination with the baseline CBF measurement yields an estimate of baseline CMRO_2_; and FLAIR-GESSE to measure tissue R_2_*′* to estimate the scaling parameter needed for calculating the change in CMRO_2_ in response to a stimulus with the calibrated BOLD method. Here we describe results for a study sample of 17 subjects (8 female, mean age=25.3 years, range 21-31 years). The primary findings were that OEF values measured with the VSEAN method were in good agreement with previous PET findings, while estimates of the dynamic change in CMRO_2_ in response to a visual stimulus were in good agreement between the traditional hypercapnia calibration and calibration based on R_2_*′*. These results support the potential of gas-free methods for quantitative physiological measurements.

**Synopsis:** We tested noninvasive methods to measure absolute oxygen metabolism (CMRO_2_) in both baseline and activation states without the use of special gases: VSEAN to measure baseline O_2_ extraction fraction (OEF), and FLAIR-GESSE to measure R_2_*′* to estimate the scaling parameter *M*. Primary findings were: CMRO_2_ changes to visual stimulation derived from R_2_*′* were similar to estimates based on hypercapnia-derived *M*; OEF values were in good agreement with previous PET findings; and, variation of baseline CBF/CMRO_2_ coupling across subjects does not follow activation coupling, suggesting different mechanisms may be involved. These results support the potential of gas-free methods for quantitative physiological measurements.

**Purpose:** To demonstrate the potential for two non-invasive techniques, VSEAN and FLAIR-GESSE, for absolute measurements of CMRO_2_ during both baseline and activation states.

## 1. Introduction

The blood-oxygenation-level dependent (BOLD) signal has been used extensively for mapping changes of neural activity noninvasively in the human brain. However, the physiological complexity of the BOLD signal makes interpretation of BOLD data challenging, particularly in studies of development and disease. Quantitative fMRI methods, designed to measure the two physiological variables underlying the BOLD response—cerebral blood flow (CBF) and the cerebral metabolic rate of oxygen (CMRO_2_)—may paint a more complete picture of neural physiology than BOLD alone. CMRO_2_ reflects the energy cost of neural activity and, when paired with CBF, may potentially provide information on the activity of specific neural populations (Buxton et al., 2014; Uhlirova et al., 2016b, 2016a). In addition, quantitative physiological measurements have the potential to expand clinical applications of fMRI, such as yielding fundamentally different conclusions about underlying physiology than possible with BOLD alone. The primary goal of technological development in this area is to measure both baseline values and activation changes in CBF and CMRO_2_ using quantitative physiological units. Quantitative and noninvasive measurements of CBF using arterial spin labeling (ASL) methods are now well established (Alsop et al., 2015). ASL allows for quantitative measurement of CBF across multiple image slices within a few seconds, depending on the technique and imaging volume. Given this short acquisition time, serial ASL images acquired over several minutes can either be averaged together to improve signal-to-noise ratio for average CBF measurement, or used as a dynamic time series to study CBF fluctuations over time (Griffeth and Buxton, 2011; Liu et al., 2019; Perthen et al., 2008).

In contrast, measuring CMRO_2_ is challenging with any technique (Buxton, 2010), and measuring baseline and dynamic change must be approached with two different classes of methods. The *fractional* change in CMRO_2_ can be measured with a calibrated BOLD approach based on simultaneous dynamic measurements of ASL and BOLD signals (Davis et al., 1998). This approach requires an additional calibration experiment to determine a scaling parameter *M* in the model of the BOLD signal. *M* depends on the amount of deoxyhemoglobin in the baseline state of the individual, and directly scales the BOLD signal measured for that individual for given changes in CBF and CMRO_2_. With this measured value of *M*, the CBF and BOLD responses to neural activation are analyzed with the same model of the BOLD signal to estimate the fractional change in CMRO_2_. In the classic calibrated-BOLD method, *M* is calculated on a voxel-wise or region-of-interest (ROI) basis from measured CBF and BOLD responses to a hypercapnia challenge (Davis et al., 1998), with the assumption that the elevated CO_2_ produces a change in CBF with no change in CMRO_2_. *Baseline* CMRO_2_ is approached by measuring the baseline oxygen extraction fraction (OEF), and several techniques have been proposed based on different ways in which the oxygenation of blood affects the MR signal. A current technique uses measurements of the BOLD response to breathing a gas with elevated O_2_, in conjunction with the measurements of the response to CO_2_ (Bulte et al., 2012; Gauthier and Hoge, 2013; Merola et al., 2018; Wise et al., 2013).

This need to make measurements while the subject breathes controlled gas mixtures in the multi-gas approach makes quantitative fMRI a complicated procedure that limits wider application beyond research settings. In addition, wearing gas masks or breathing higher concentrations of CO_2_ and O_2_ would not be possible in some patient populations. To eliminate this barrier to entry into clinical applications, and for more widespread application of quantitative fMRI methods in basic brain studies, we evaluated two methods for measuring baseline and activation changes in CMRO_2_ that do not require the subject to breathe special gas mixtures. The calibration information to estimate *M* is based on measurement of tissue R_2_*′*, essentially the component of transverse magnetization relaxation that can be reversed with a spin echo. Recent theoretical work (Blockley et al., 2015; Blockley and Stone, 2016), based on earlier seminal work of Haacke and Yablonskiy (Yablonskiy, 1998; Yablonskiy and Haacke, 1994), showed that R_2_*′*, like *M*, essentially depends on the total deoxyhemoglobin content of tissue. In this study, R_2_*′* was measured with a modified FLuid Attenuated Inversion Recovery Gradient Echo Sampling of Spin Echo sequence (FLAIR-GESSE) method (Simon et al., 2016). The baseline CMRO_2_ measurement was done with a technique called Velocity-Selective Excitation and Arterial Nulling (VSEAN) (Guo and Wong, 2012) that isolates the signal of local venous blood and measures its relaxation rate R_2_. Because R_2_ depends primarily on the O_2_ saturation of hemoglobin, applying an appropriate calibration curve provides a measurement of the local O_2_ extraction fraction (OEF), and together with a baseline CBF measurement from ASL provides a measurement of baseline CMRO_2_. In this study, we evaluated the combination of these two gas-free techniques to make absolute baseline and activation CMRO_2_ measurements as an initial step toward more widespread applications of quantitative fMRI methods.

## 2. Methods

### 2.1 Subjects

Twenty-one healthy adults were recruited for the study. Inspired and end-tidal O_2_ and CO_2_ were monitored for all subjects throughout the run. Two of the 21 subjects were eliminated from the analyses because inspired and end-tidal O_2_ and CO_2_ measurements revealed leaks in the tubing or non-rebreathing facemask. Two other subjects exhibited no BOLD or flow change to the visual stimulus administration, demonstrating that the visual cortex was not well targeted. Thus, the study sample included 17 subjects (8 female, mean age=25.3 years, range 21-31 years). The study was approved by the Human Research Protections Program of the University of California, San Diego; written informed consent was obtained from all subjects. Subjects were remunerated for their participation.

### 2.2 Gas administration

Subjects were equipped with a non-rebreathing facemask (Hans Rudolph, KS, USA). The inspiratory port of the mask was connected to a large gas-tight balloon (VacuMed, CA, USA), pre-filled with a pre-mixed hypercapnia gas mixture of 5% CO_2_, 21% O_2_, balance N_2_ (Airgas-West, CA, USA). The tubing (VacuMed, CA, USA) was disconnected to allow the subject to breathe normal room air in the normocapnic condition. End-tidal and inspired O_2_ and CO_2_ were monitored using a Perkin Elmer 1100 medical gas spectrometer (Perkin Elmer, Waltham, MA).

### 2.3 Imaging

#### 2.3.1 BOLD-ASL

A dual-echo gradient echo spiral PICORE QUIPSS II ASL acquisition (Wong et al., 1998) was used to acquire simultaneous BOLD and ASL dynamic images on a General Electric (GE) Discovery MR750 3.0T scanner. Seven axial slices (5 mm thick/1 mm gap) covering the occipital cortex, centered around the calcarine sulcus, were obtained with TR=2500ms, TI1=700ms, TI2=1750ms, TE=3.3/30ms, 90° flip angle, FOV 256 mm × 256 mm, and matrix 64×64. Field maps with the same slice prescription were acquired to correct distortions in the spiral acquisition due to magnetic field inhomogeneity (Noll et al., 2005). Physiological monitoring was performed throughout the scan session using a pulse oximeter for cardiac cycle monitoring and respiratory bellows for respiratory dynamics (GE MR750 built-in).

#### 2.3.2 VSEAN

The baseline CMRO_2_ measurement was done with Velocity-Selective Excitation and Arterial Nulling (VSEAN) (Guo and Wong, 2012) that isolates the signal of local venous blood and measures its relaxation rate R_2_. Because R_2_ depends primarily on the O_2_ saturation of hemoglobin, this provides a measurement of the local O_2_ extraction fraction (OEF). This method differentiates the signal of slowly moving blood from that of static tissue by velocity selective excitation, i.e., selectively generating signal only from slow moving spins in arterioles and venules. An arterial-nulling preparation module isolates the venous blood signal, measured with multiple T_2_-preparations of different effective TEs (eTEs) (Guo and Wong, 2012). The sole center slice of the BOLD-ASL prescription was used as the single 10mm VSEAN slab. Details of the acquisition are as follows:

##### Arterial nulling

A slab-selective inversion pulse (arterial inversion slab thickness = 150mm) inverted a bolus of arterial blood below the imaging plane, with an inversion/delay time TI=1150ms to allow the inverted bolus of arterial blood to arrive at the imaging plane at the null point of the arterial blood’s longitudinal magnetization during image acquisition. The spins of the static tissue and venous blood in the imaging slice were unperturbed, giving a strong venous signal.

##### Velocity-Selective Excitation (VSE) for flow signal separation

Two VSE pulse modules (one BIR4-based pulse train with velocity-sensitive gradient pulses, the second built into the image acquisition using only velocity-sensitive gradient pulses) excited and separated slow moving spins without generating signal from static or extremely slow flowing spins.

##### T_2_ measurement and oxygenation estimation

T_2_ preparation module (eTEs of 25/50/75ms with incremental gaps between RF pulses, built into the first BIR-4 based VSE module) measured T_2_. The T_2_ values were translated to blood oxygen saturation of hemoglobin Y via a T_2_-Y calibration curve. We used the calibration curve from Zhao and colleagues based on a bovine blood sample measured *in vitro* at 3 Tesla (Zhao et al., 2007), which used a similar T_2_ measurement setup as in this study. The equation for the curve used in this analysis is 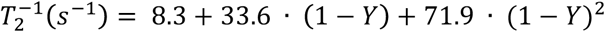 (Guo and Wong, 2012). The OEF was then taken as 1-Y, based on the assumptions that arterial hemoglobin was fully saturated with oxygen and dissolved O_2_ was negligible compared to hemoglobin-bound O_2_.

##### Other pulse sequence parameters

FOV = 256 mm × 256 mm, matrix = 64 × 64, single-slice spin echo with spatial-spectral excitation, two slice-selective hyperbolic secant refocusing pulses, single-shot spiral readout, TR/TE=3s/28ms, v_e_=2 cm/s in slice-selective direction, slice-selective post-saturation pulses, 81 acquisitions preceded by two dummy scans and including six “cos” modulated reference scans (Guo and Wong, 2012). The total scan time was 4:03.

#### 2.3.3 FLAIR-GESSE

We measured R_2_*′* with a FLuid Attenuated Inversion Recovery Gradient Echo Sampling of Spin Echo (FLAIR-GESSE) (Simon et al., 2016) technique. FLAIR-GESSE measures a series of gradient echo samples around a spin echo following Yablonskiy and Haacke’s GESSE method with two main modifications (Simon et al., 2016). First, a CSF-nulling FLAIR module is added to minimize CSF contamination of R_2_*′* measurements. Our previous work has found that CSF-nulling improves the stability of R_2_*′* measurements (Simon et al., 2016). Second, the two sides of the echo are sampled with two acquisitions; each has a different spin echo time so that the acquired data are at the same absolute time after excitation. This helps correct for problems with multiple T_2_ values within a voxel. Since R_2_*′* is sensitive to large-scale field inhomogeneity in addition to sub-voxel inhomogeneity due to deoxyhemoglobin, field offsets are acquired using phase images from the FLAIR-GESSE acquisition that are then used to calculate the large-scale field inhomogeneity and remove their contribution to the R_2_*′* estimate (Dickson et al., 2010; Simon et al., 2016). The center of each slice from the BOLD-ASL prescription was used, though for FLAIR-GESSE the slices were 2 mm thick (to minimize the effect of through-plane field gradients), with a gap of 4 mm. The pulse sequence parameters included two pairs of GESSE image series, one as an early spin echo series (63.6 ms after excitation) and one as a late spin echo series (83.6 ms after excitation), collected separately. For each spin echo series, 64 samples of each decay curve were collected asymmetrically. CSF nulling was chosen for middle slices, with a TR of 3.5 s, inversion time TI of 1.16 s, matrix = 64 × 64, and FOV = 256 mm × 256 mm. Each image slice was acquired in ascending order with a spacing of 110 ms. Each of the two spin echo series was 3:58 in duration. R_2_*′* was calculated by the method described in Simon et al., 2016.

#### 2.3.4 Calibration and reference scans

A cerebral spinal fluid (CSF) reference scan using a single-echo spiral EPI acquisition (TE = 3.3ms) was obtained for CBF quantification (Chalela et al., 2000; Liu et al., 2004; Perthen et al., 2008). A minimum contrast scan (eight-shot spiral acquisition, TE=11ms, TR=2000ms) was made to correct for transmit and receive coil inhomogeneities (Wang et al., 2005). These reference and calibration scans used the same prescription as the ASL acquisition. A high resolution T_1_-weighted anatomical scan (FSPGR) was acquired for image registration and segmentation.

### 2.4 Stimulus paradigm

The visual stimulus task used to elicit neural activity in the occipital lobe (V1) was a black and white flickering radial checkerboard (6 Hz light-dark reversal frequency) with contrast and luminance as described in previous work (Simon et al., 2016). The central region was maintained at iso-luminant gray with visual angle ∼1.5°. The stimulus was presented using MATLAB (2014a, The MathWorks, MA, USA) with the Psychophysics Toolbox extensions (Brainard, 1997; Pelli, 1997). The subject viewed the stimulus, projected onto a screen, through a mirror set atop the head coil.

### 2.5 Experimental setup

BOLD and ASL responses were measured during a baseline task and to a visual stimulus in normocapnia. A hypercapnia calibration measurement and separate VSEAN and FLAIR-GESSE acquisitions in normocapnia were performed with the baseline task. The baseline state for all acquisitions performed in this experiment was a 1-back task (Kirchner, 1958) projected on a cross at the center of the screen. Subjects fixated at this center cross and performed the 1-back task that presented single digits sequentially at random in 1-second intervals. The subjects were instructed to press a button on a response box each time they observed a number repeated sequentially. As this served as the baseline state, the task was performed continuously throughout each acquisition, with the ON or OFF states representing the presence of a visual stimulus, which was the flickering checkerboard described above filling the visual field surrounding the center cross. Given the 11-minute duration of the BOLD/ASL acquisition, the 1-back task was chosen as the baseline rather than a resting state (visual fixation on a cross) to hold the subject’s attention throughout the long scan.

The BOLD/ASL acquisition consisted of an 11-min run, starting at normoxia/normocapnia with 3 blocks of 30-sec OFF/30-sec ON visual localizer at the beginning to independently determine an activated visual region of interest (ROI). The following experimental stimuli consisted of 2 blocks of 1-min OFF/1-min ON visual stimulus. The final 4 minutes of the BOLD/ASL acquisition was the calibrated BOLD experiment with a 1-min baseline period at normoxia/normocapnia, then a 3-min block of 5% CO_2_ administration (normoxia/hypercapnia). A schematic of the run is shown in Figure 1.

**Figure 1:**
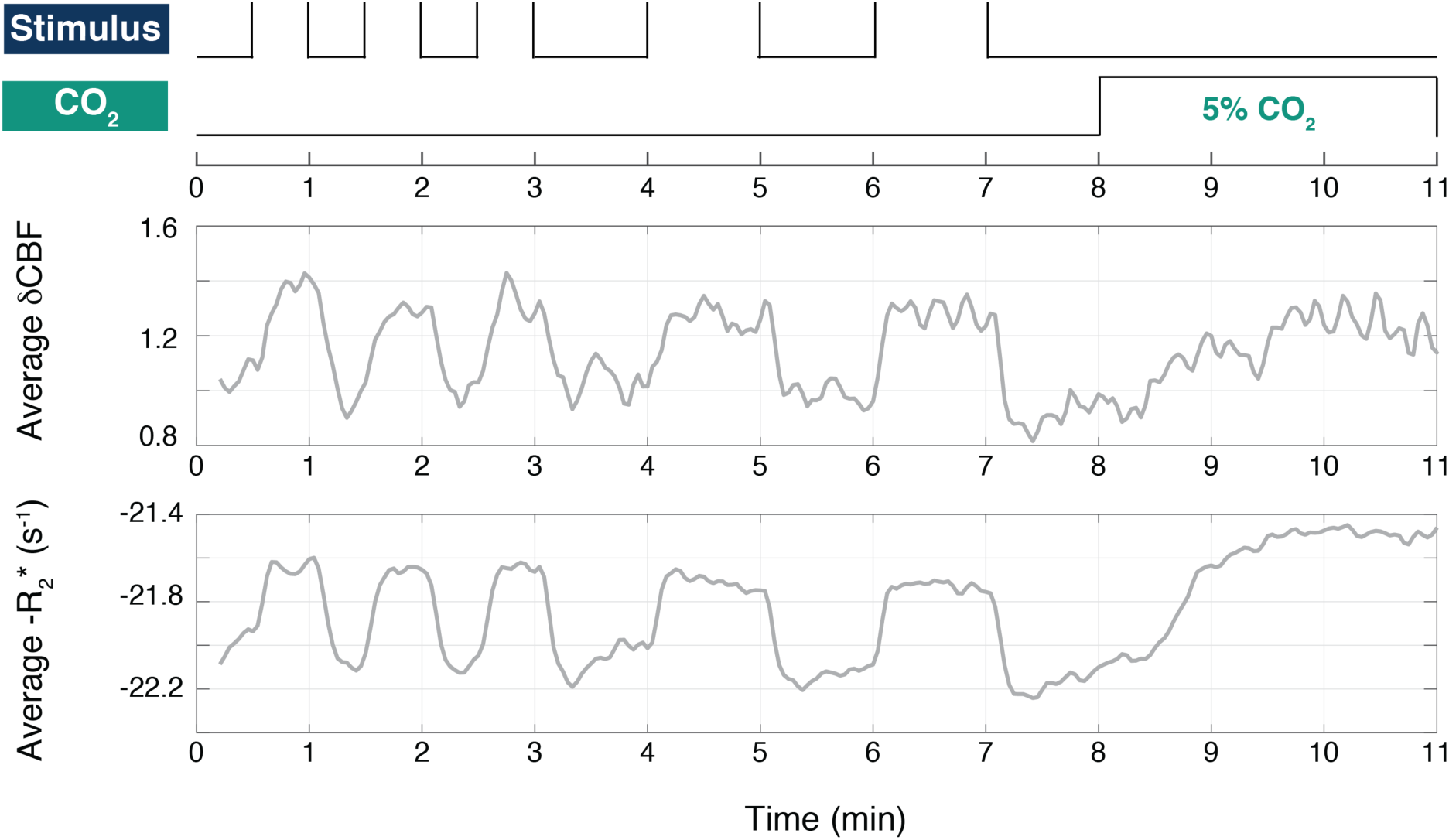
Experimental set-up, CBF and R_2_* data. Top panel shows experimental set-up of PICORE QUIPSS II BOLD/ASL acquisition with visual stimuli of a 6 Hz flickering checkerboard (“Stimulus”) and 5% CO_2_ administration. Middle and bottom panels illustrate average CBF and BOLD R_2_* curves across 17 subjects. The CBF is normalized to the baseline CBF, and thus is dimensionless. The BOLD R_2_* curve is plotted as –R_2_* for more visibly intuitive purposes.

The FLAIR-GESSE and VSEAN acquisitions as described above were all acquired at the baseline state (1-back task with no visual stimulus). Calibration and reference scans were acquired pre- and post-experimental acquisitions.

### 2.6 Data preprocessing

#### 2.6.1 Arterial Spin Labeling (ASL) images

Raw ASL images were first corrected for inhomogeneity in the magnetic field using the field map acquisition (Noll et al., 2005). The functional scan was motion corrected and registered to one image in the 11-minute run using AFNI software (Cox, 1996). The first four images of the run were discarded to allow the signal to reach steady state. Minimum contrast images were used to correct ASL data for coil sensitivity inhomogeneity (Wang et al., 2005). Applying surround-subtraction to the raw first-echo ASL images produced CBF-weighted images with minimal contamination from BOLD (Liu and Wong, 2005), and these were converted to physiological units (ml/100ml/min) using CSF as a reference (Chalela et al., 2000). A gray matter mask was determined from the absolute CBF, in which all voxels with CBF value greater than twice the mean CBF, taken as the average across all voxels, were included as part of the GM mask for each subject. Thus, the masks are generated from the BOLD/ASL acquisition, circumventing the need for anatomical registration to the functional data that could potentially distort voxels and thus the measurements. Empirically, we have found this threshold to give a good correspondence with GM masks generated from T_1_-weighted anatomical images when the separate acquisitions are registered. For the visual ROI determination in 2.6.3, the gray matter mask was limited to the posterior third of the brain via a binary wedge-shaped mask covering the posterior third.

#### 2.6.2. BOLD R_2_*-weighted images

BOLD R_2_*-weighted images were calculated from surround averaging of the first and second echo ASL images (Liu and Wong, 2005) to enable calculation of R_2_* for each time point as previously reported (Liu et al., 2019). For each time series, after ROI selection (Section 2.6.3), the mean R_2_* was subtracted to form a time series ΔR_2_*(t), which removed systematic effects of drift that scale identically with both echoes; ΔR_2_* was then converted back to a *δ*BOLD signal without the drift effects (Liu et al., 2019).

#### 2.6.3 Regions of interest (ROIs)

Visual ROIs were generated using the ASL and BOLD responses to the 3-min visual functional localizer exclusively, then limited to the center slice that was used for VSEAN. A general linear model approach, described by Perthen et al. (Perthen et al., 2008), was used for ROI selection. The pattern of the stimulus was convolved with a gamma density function to produce a stimulus regressor (Boynton et al., 1996). A constant and a linear term were used as nuisance regressors. A mask containing only gray matter voxels, as determined in 2.6.1, in the posterior third of the brain was used to further restrict the ROI selection. Voxels exhibiting both CBF and BOLD activations were detected using an overall significance threshold of p=0.05 and minimum cluster size of 2. Thus, the final visual ROI for each subject consisted of a single mask of voxels exhibiting independent activation in both CBF and BOLD to the visual stimulus, restricted to gray matter in the posterior third of the brain.

#### 2.6.4. Average CBF and BOLD changes to stimuli

Average CBF and R_2_* from BOLD (Section 2.6.2) time courses for the ROI were calculated for each subject, producing one-dimensional BOLD and CBF time courses. The resulting time series were normalized to an average baseline value, defined as the mean of the 15 seconds of rest before the first normocapnia visual stimulus (at the 4-min mark), the 15 seconds of rest before the second normocapnia visual stimulus (at the 6-min mark), and the 15 seconds of rest before the CO_2_ stimulus onset (at the 8-min mark). These normalized values were used for all subsequent analyses. To allow time for the subjects’ inspired and end-tidal gas levels to stabilize and equilibrate, data from the first minute of the hypercapnia block were not used subsequently. To derive single estimates of CBF and BOLD responses for each subject, the two visual stimulus blocks were first averaged.

### 2.7 Calibrated BOLD and FLAIR-GESSE analysis

In the classic calibrated-BOLD method (Davis et al., 1998) to measure the fractional change in CMRO_2_ with activation, the BOLD signal is modeled as:

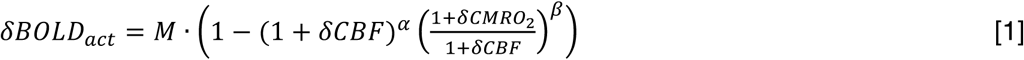

where the prefix “*δ*” indicates the change in the variable normalized to the baseline value. The scaling parameter *M* is approximately proportional to the total deoxyhemoglobin content of tissue, which could vary across brain regions and subjects. The parameters *α* and *β* were originally introduced to describe specific physical effects related to the change in venous blood volume and nonlinearities of the magnetic susceptibility effects, respectively (Davis et al., 1998). However, that original derivation left out several factors affecting the BOLD signal, and later modeling studies including these factors found that the mathematical form of Eq. [1] is still accurate, but the parameters *α* and *β* are now thought of more as fitting parameters rather than reflecting their original physical meanings (Buxton, 2013; Gagnon et al., 2016, 2015; Griffeth and Buxton, 2011; Merola et al., 2016). The standard approach to estimate *M* is to measure CBF and BOLD responses to hypercapnia, and analyze the data with Eq. [1] and the assumptions: 1) there is no CMRO_2_ change during hypercapnia; and 2) particular values of *α* and *β*, typically *α*=0.2 and *β*=1.3 (Chen and Pike, 2009; Griffeth et al., 2013; Mark et al., 2011). With these assumptions, the scaling factor *M* is calculated from the hypercapnia data. The activation data are then analyzed with the estimated value of *M* and the same assumed values for *α* and *β* to estimate the fractional CMRO_2_ change to the stimulus.

In the current study, we compared the classic hypercapnia estimate of *M* with a measurement of R_2_*′*, approximately the difference in transverse relaxation rates measured in gradient echo and spin echo experiments: R_2_*′* ≅ R_2_* - R_2_. The parameter R_2_*′* is sensitive to magnetic field inhomogeneities due to 1) deoxygenated blood vessels, proportional to voxel baseline deoxyhemoglobin concentration, like *M*, and 2) large-scale inhomogeneities of the head. The inhomogeneity correction was done as in Simon et al., 2016, and the remaining R_2_*′* value was interpreted as reflecting deoxygenated blood vessels. Neglecting some of the complexities of tissue relaxation (see Discussion for applicability and limitations), we would expect *M* ≅ *M′* = TE • R_2_*′* (Blockley et al., 2015).

FLAIR-GESSE data were analyzed per Simon, et al. 2016. The raw GESSE data were first averaged across the visual ROI before R_2_ and R_2_*′* values were fitted to the averaged data. Magnetic field inhomogeneities were corrected per Dickson et al. (Dickson et al., 2010). *M′* was calculated from R_2_*′* values through *M′* ≅ TE • R_2_*′*. Fractional change in CMRO_2_ (*δ*CMRO_2_) to the visual stimulus for each individual was calculated per Eq. [1] (*α*=0.2, *β*=1.3) using the individual *M′* value for each subject, as well as with the group mean *M′*, to compare estimates of *δ*CMRO_2_ using individual *M′* versus group mean *M′*. Meanwhile, the hypercapnia response BOLD data were also substituted into Eq. [1] to yield an estimate of *M* for each subject, calculated for *α*=0.2, *β*=1.3. *δ*CMRO_2_ to the visual stimulus for each individual was then calculated per Eq. [1], (*α*=0.2, *β*=1.3) using the individual *M* value for each subject, as well as with the group mean *M*, to compare estimates of *δ*CMRO_2_ using individual *M* versus group mean *M*. These values were compared to those using *M′*.

### 2.8 VSEAN and baseline CMRO_2_ analysis

VSEAN data were analyzed per Guo and Wong (2012). Raw multi-echo data were averaged over the visual ROIs first, then used to fit a T_2_ value representative of the entire ROI. The T_2_ was then converted into blood oxygenation (Y) levels via T_2_-Y calibration curve at 3T (Zhao et al., 2007). OEF was calculated from venous oxygenation (OEF = 1 – venous oxygenation). Baseline CMRO_2_ was calculated from

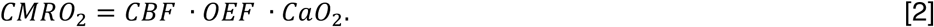

The equation to determine total arterial oxygen concentration is: CaO_2_ = 1.36(Hb)(S_a_O_2_/100) + 0.0031(P_a_O_2_), with units as follows: CaO_2_ in ml/dL, 1.36 ml O_2_/1 g Hb, Hb in g/dL, S_a_O_2_ in %, 0.0031 g/dl/mmHg, and P_a_O_2_ in mmHg (Davenport, 1974). Hemoglobin concentration was assumed according to mean healthy ranges set by UC San Diego Health (Fraser and Haldeman-Englert, 2017); for females, [Hb] = 14g/dL; males, [Hb]=15.7g/dL. The age of each subject was also factored into CaO_2_ determination through the estimate of P_a_O_2_ (Estimated normal P_a_O_2_ ≅ 100 mmHg – (0.3) age in years) (Crawford and Adesanya, 2010; Jurado and Walker, 1969). CBF is expressed in ml/100ml/min, and CaO_2_ from this equation is divided by 100 to give CMRO_2_ units in ml/100ml/min. CMRO_2_ in ml/100ml/min is then converted to mM/min (1 mL O_2_ at 310 K, 1 atm = 0.03933 mmol O_2_) (Davenport, 1974). For reference, 1 mM/min = 2.54 ml O_2_/100 ml/min = 2.42 ml O_2_/100g/min = 100 umol/100 ml/min = 95 umol/100g/min, with assumed density of blood = 1.05 g/mL at 310 K (Trudnowski and Rico, 1974).

### 2.9 Voxel-wise analysis

To compare the quality of VSEAN and FLAIR-GESSE data on a voxel-wise basis to ROI-based analyses, maps of OEF and R_2_*′* values were made for each voxel in the visual ROI. The masks were then further limited to encompass voxels that survived both to create dually-restricted (VSEAN and FLAIR-GESSE restricted) masks for the ROI. The median, mean, and standard deviation of the voxel-wise OEF and R_2_*′* estimates were calculated for each subject, then averaged across all 17 subjects.

### 2.10 Data and code availability

The data that support the findings of this study are available from the corresponding author, RBB, upon reasonable request. The data and code sharing adopted by the authors comply with requirements by the National Institutes of Health and the University of California, San Diego, and comply with institutional ethics approval.

## 3. Results

The following results present mean findings from the visual stimulus ROI (section 2.6.3). Voxelwise findings from the visual ROI and gray matter are described in the Supplementary Materials. Those findings are not significantly different from the mean OEF and R_2_*′* calculated from the ROI-based method described here.

### 3.1 BOLD/ASL results

Average CBF and BOLD R_2_* curves from 17 subjects across the entire 11-min acquisition are plotted in Figure 2. The average fractional change in CBF and BOLD to the visual stimulus and CO_2_ challenge for each subject are plotted in Figure 2A. The average responses of the cohort are summarized in Figure 2B.

**Figure 2:**
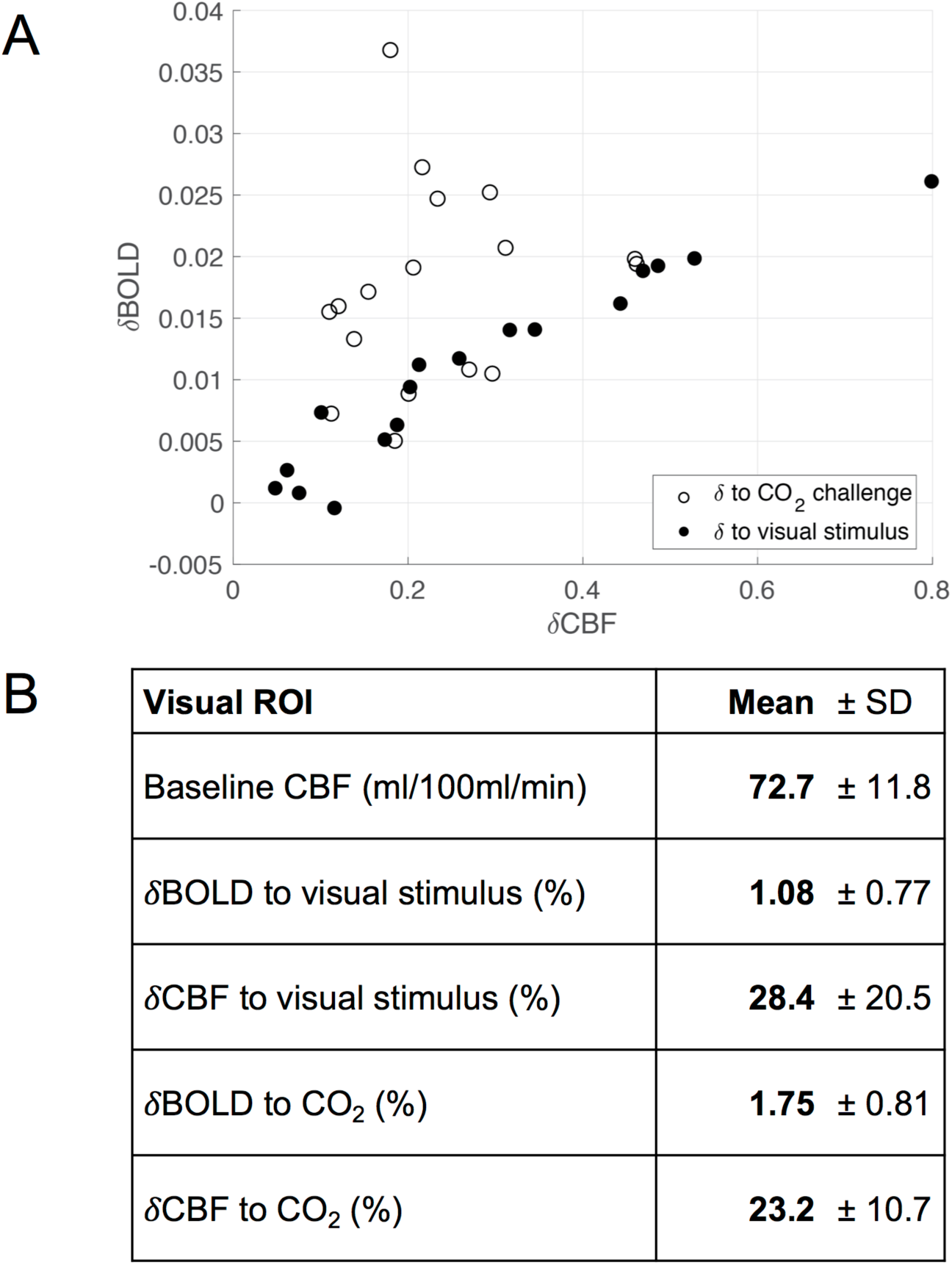
Visual activation and calibrated BOLD results. (A) Scatter plot of fractional CBF versus fractional BOLD change to the visual stimulus and CO_2_ challenge for each subject in the visual ROI only. (B) Mean and standard deviation of *δ*BOLD and *δ*CBF to visual stimulus experiment in normoxia/normocapnia across seventeen subjects, and mean changes to CO_2_ challenge in the baseline state, across all subjects.

### 3.2 VSEAN results

Figure 3 shows the OEF measurements made using VSEAN within the visual ROI plotted against the baseline CBF (Fig. 3A), absolute ΔCBF to CO_2_ (Fig. 3B), and absolute ΔCBF to visual activation (Fig. 3C). Pearson correlation analyses found no significant association with baseline OEF in any of these conditions (Baseline CBF: *r* = -.188, *p*=0.470, Cohen’s *d*=0.38; Absolute ΔCBF to CO_2_: *r* = -.248, *p*=0.336, *d*=0.51; ΔCBF to visual activation: *r* = -.054, *p*=0.837, *d*=0.11). The findings in Figs. 3A-C suggest possible sex differences in the CBF measures. Figure 3D summarizes OEF and CBF values of the entire sample, and separately for males and females. A two-way repeated-measures ANOVA revealed a significant main effect of sex (*F*_1,15_=7.103, *p*=.018, *d*=1.38) with no significant sex by measure interaction (*F*_3,45_=2.096, *p*=.114, *d*=0.75). Post-hoc analyses showed females had greater baseline CBF and absolute ΔCBF to CO_2_ than males (*t*_*15*_ = 3.090 & 2.843, Bonferonni corrected *p* = .028 & .048, *d* = 1.51 & 1.35, respectively). No significant sex differences were detected in OEF or absolute ΔCBF to visual activation (*t*s < 0.25, *p*s > .810, *d*s < 0.12). Similar effects have been reported previously in work studying regional differences in CBF by sex (Esposito et al., 1996; Gur and Gur, 1990; Rodriguez et al., 1988).

**Figure 3:**
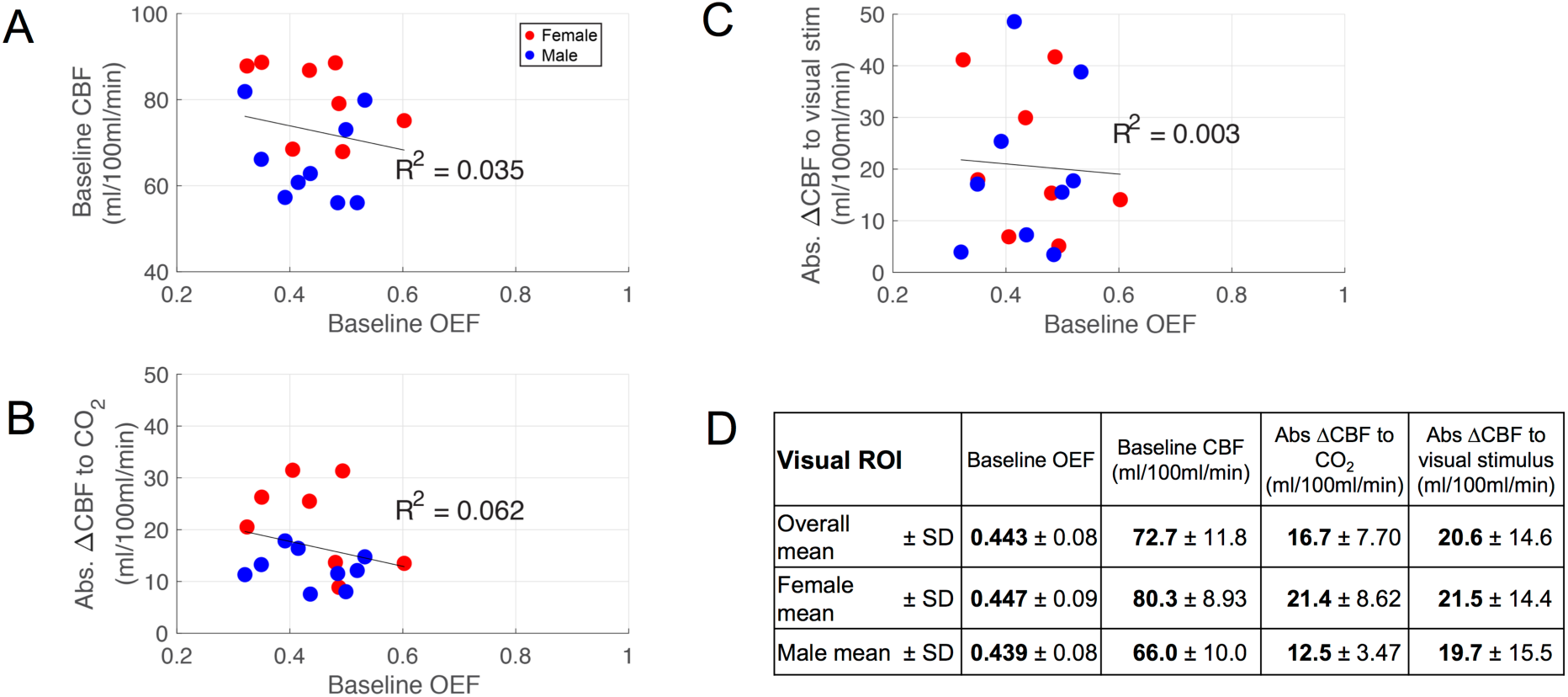
VSEAN results. (A) Independent VSEAN and ASL methods show variation of measured baseline OEF and CBF across subjects in the visually stimulated ROI. (B) Variation of OEF to absolute ΔCBF to CO_2_ are plotted for ROI, and (C) variation across subjects versus absolute ΔCBF to visual stimulus in the visual ROI. (D) Table of baseline OEF, baseline CBF, absolute ΔCBF to CO_2_, and absolute ΔCBF to visual stimulus in visual ROI. There was no correlation between OEF and baseline CBF or to any absolute ΔCBF. There was a significant and large effect size difference in baseline CBF between male (blue) and female (red) (p=0.007, Cohen’s *d* = 1.5).

### 3.3 FLAIR-GESSE and M calibration results

Mean measured R_2_*′* across 17 subjects in the visual ROI was 3.60 s^-1^, with a standard deviation of 1.39 s^-1^, giving a mean value of *M′*=0.108 +/- 0.04 (Figure 4B). The calibrated BOLD analysis of the hypercapnia data yielded an average estimate of *M* = 0.095 +/- 0.05 (Figure 4B). Figure 4A shows a plot of the individual data for *M′* against the value of *M* calculated for *α*=0.2 and *β*=1.3. We also compared *M* and *M′* with OEF for each subject and found no statistically significant correlation (see Supplementary Materials for details). *δ*CMRO_2_ calculated using individual and group mean *M* and *M′* are reported in Figure 4C. While individual and group mean *M′* values yielded similar *δ*CMRO_2_, individual *M* from hypercapnia yielded significantly smaller mean *δ*CMRO_2_ than that of the group mean (p=0.049).

**Figure 4:**
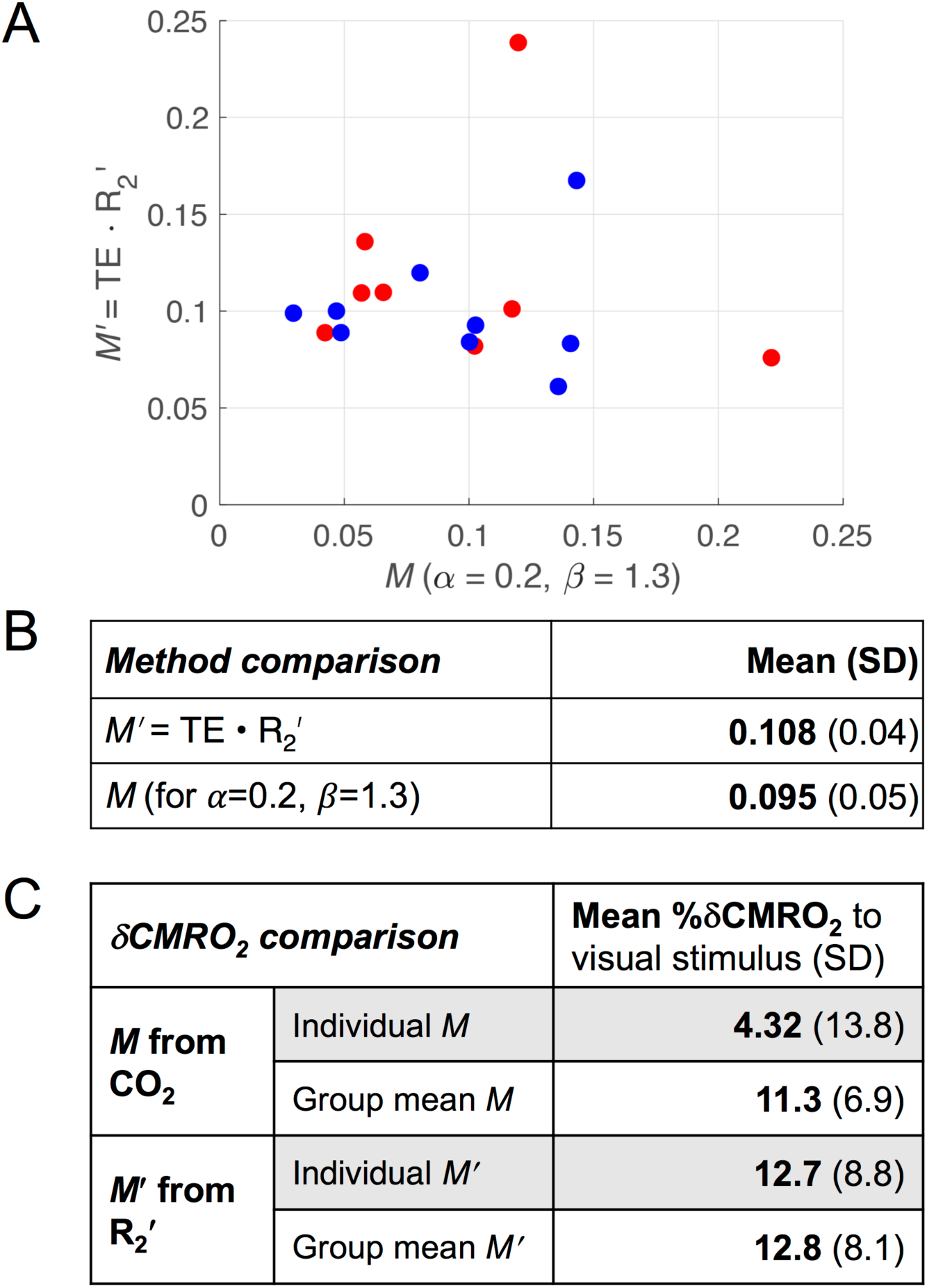
FLAIR-GESSE versus calibrated BOLD results. (A) *M* from calibrated-BOLD experiment with *α*=0.2, *β*=1.3, and *M′* from FLAIR-GESSE’s R_2_*′* for each of 17 subjects in the visual ROI. Scatterplot demonstrates range and variance of measurements of *M* from the two methods. (B) Average *M′* was calculated from mean measured R_2_*′* value across 17 subjects using FLAIR-GESSE of 3.60 +/- 1.39 s^-1^, with TE = 0.03 s. *M* values were calculated from calibrated BOLD CO_2_ experiment. (C) Fractional *δ*CMRO_2_ to visual stimulus calculated using Eq. [1] and *δ*CBF and *δ*BOLD to the visual stimulus, as well as the *M* and *M′* values. Two calculations were performed, one using each subject’s individual *M* or *M′* value to calculate individual *δ*CMRO_2_, then averaged across all subjects; the other method utilized the group mean *M* or *M′* to calculate individual *δ*CMRO_2_, then averaged across the subjects.

### 3.4 Absolute CMRO_2_ and CBF measurements

The measured data provided the essential information needed to make estimates of absolute CBF and CMRO_2_ in the baseline state and absolute change in CBF and CMRO_2_ to the visual stimulus, thus yielding a complete quantitative picture of underlying physiology. These relationships are shown in Figure 5. The average baseline CMRO_2_, determined using Eq. [2] and OEF=0.44 from VSEAN, was calculated to be 2.58 +/- 0.53 mM/min, equivalent to approximately 6.56 ml/100ml/min = 6.24 ml/100g/min = 258 umol/100ml/min = 245.1 μmol/100g/min (see section 2.8 for conversion). The table in Figure 5E shows the mean measured absolute change in CMRO_2_ to the stimulus using *M′* values for each subject calculated from R_2_*′*, based on *α*=0.2 and *β*=1.3. Because this estimate depends on the estimate of the fractional CMRO_2_ change, it also depends on the assumed value of *β*. The absolute ΔCMRO_2_ to a visual stimulus was estimated to be 0.21 mM/min for *β*=1.3. Figure 5 shows the spread of individual measurements for these parameters. Importantly, in Figures 5A and 5B, the *x* and *y* axes are not independent measures as the value of baseline CMRO_2_ was calculated from baseline CBF.

**Figure 5:**
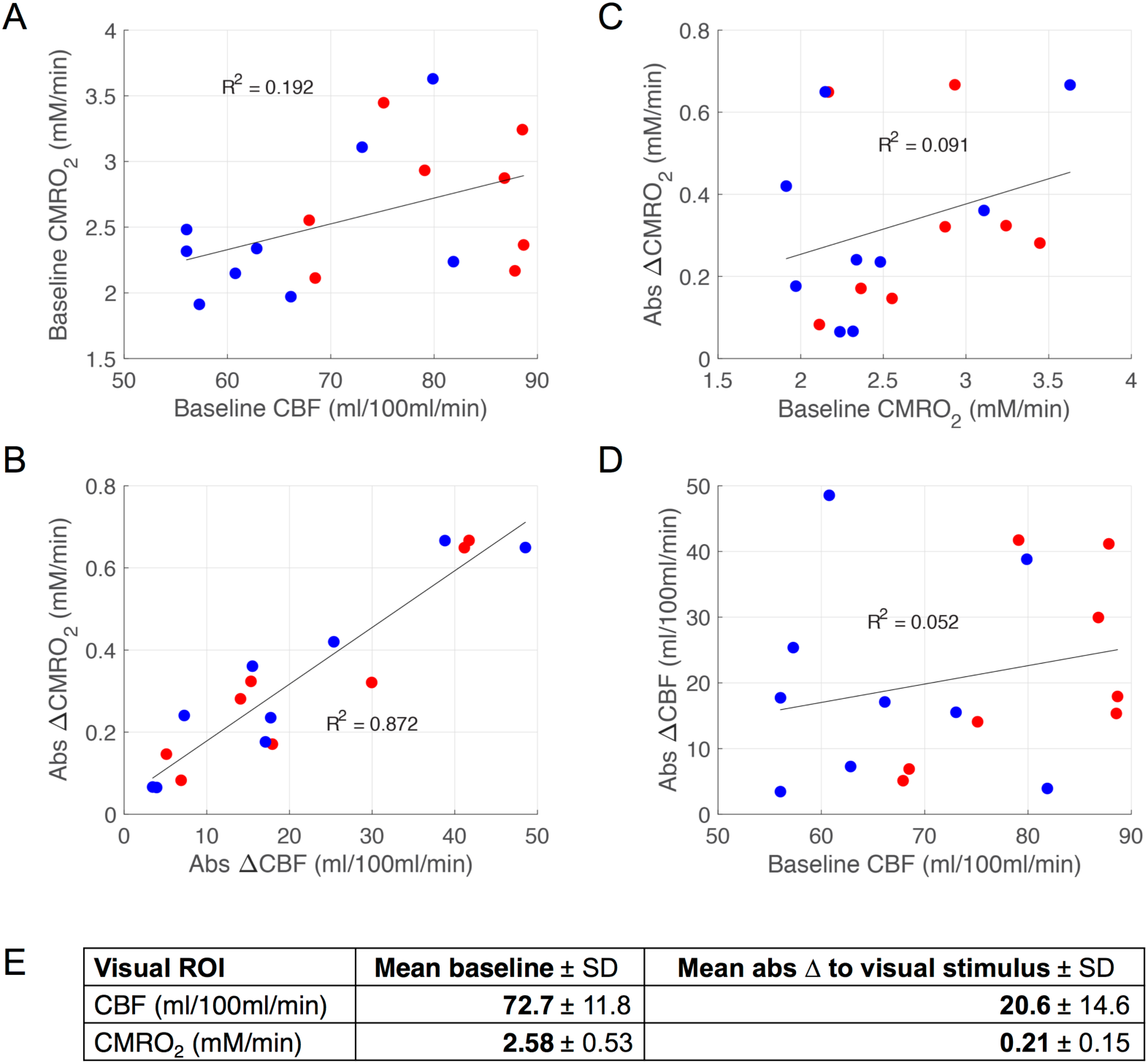
Baseline and absolute changes in CMRO_2_ and CBF across subjects in visual ROI. Absolute calculations of CMRO_2_ at baseline and to a visual stimulus non-invasively determined using OEF and *M* calibration data acquired from VSEAN and FLAIR-GESSE, with *α*=0.2 and *β*=1.3. (A) Absolute CBF and CMRO_2_ in the baseline state (slope=0.020, *r*=0.438, p=0.079, Cohen’s *d* = 0.97). (B) Absolute CBF and CMRO_2_ changes in response to the visual stimulus (slope=0.014, *r*=0.934, *p*<0.00001, *d* = 5.23). Note that x- and y-axes are not independent in A or B as CMRO_2_ estimates are calculated from CBF. (C) Absolute ΔCMRO_2_ change to visual stimulus versus baseline CMRO_2_. Change in CMRO_2_, based on fractional change of BOLD and CBF to visual stimulus, was calculated independently of baseline CMRO_2_, which is dependent on baseline CBF and an independent VSEAN measurement of OEF. The change in CMRO_2_ was calculated using *M′* from individual R_2_*′* (slope=0.091, *r*=0.302, *p*=0.079, *d* = 0.0.63). (D) Absolute ΔCBF change to visual stimulus versus baseline CBF. Measurements are independent; baseline CBF was calculated based on the CSF calibration scan, while ΔCBF was estimated from ASL data with visual stimulus onset (slope=0.052, *r*=0.228, *p*=0.379, *d* = 0.47). (E) Mean baseline and absolute change estimates for CMRO_2_ and CBF.

## 4. Discussion

This work explored the feasibility of making absolute measurements of CMRO_2_ and CBF without requiring the subject to breathe special gases as a first step toward more widespread application of quantitative physiological measurements. While arterial spin labeling (ASL) techniques that do not involve gases are well developed to measure both baseline and fractional change in CBF, CMRO_2_ measurement is much more challenging and often involves administration of gases with higher concentrations of O_2_ or CO_2_. The main barrier to measuring baseline CMRO_2_ is obtaining a measurement of oxygen extraction fraction (OEF), while measuring fractional change in CMRO_2_ to a stimulus requires calibration of the blood oxygenation level dependent (BOLD) signal through the factor *M*.

The two gas-free techniques tested here present ways to measure those parameters noninvasively. Velocity-selective excitation and arterial nulling (VSEAN) is a technique that leverages relaxation effects of altered hemoglobin O_2_-saturation on the transverse relaxation rate (R_2_) of venous blood to determine regional OEF. Fluid attenuated inversion recovery – gradient echo sampling of spin echo (FLAIR-GESSE) is a method of estimating the factor *M* through measurement of R_2_*′* that reflects the amount of total deoxyhemoglobin in tissue. We applied these methods to a relatively homogeneous cohort of 17 subjects. Because of the uniformity of the study population, we expected that the physiological parameters were likely to be similar across subjects, and that the variance of the measured values across subjects was thus likely to be dominated by the intrinsic variability of the measurement methods. We took this conservative approach in interpreting the performance of these methods. Our primary findings were: 1) VSEAN yielded a mean value of baseline OEF in the visual ROI of 0.44 ± 0.08, matching previously reported measurements of extraction (Ibaraki et al., 2007; Ishii et al., 1996; Ito et al., 2004) using positron-emission tomography (PET); and 2) the estimated changes in CMRO_2_ in response to the visual stimulus were similar when calculated from *M′*, which was estimated from R_2_*′* measured with the FLAIR-GESSE method, and when calculated from *M* estimated from the hypercapnia response. The ability to obtain a complete quantitative picture of human brain activation, with absolute measures of CBF and CMRO_2_ in both baseline and activation states, would be a useful tool for basic studies of brain function and for clinical applications to characterize the effects of disease.

### 4.1 Baseline CMRO_2_ with VSEAN OEF measurement

The VSEAN method is an extension of the seminal method QUIXOTIC for measuring baseline OEF (Bolar et al., 2011), expanding on the basic idea to overcome some limitations of the earlier technique. QUIXOTIC uses a velocity-selective (VS) spin labeling technique, separating venous blood from arterial blood and static tissue. However, the sequence design causes the venous signal to be potentially contaminated by signal contribution from CSF due to diffusion attenuation and to be attenuated by a global inversion pulse. VSEAN addresses these issues by isolating the venous blood signal using a separate arterial nulling module and improves upon the signal to noise of measuring venous T_2_ by selective excitation of moving venous spins right before acquisition which allows more time for venous spins to recover, thus yielding a stronger venous signal in comparison to that of QUIXOTIC, where a global inversion pulse attenuates all signal, including venous signal.

Although VSEAN had limited testing in human subjects previously, this is the first study utilizing VSEAN on a larger sample of subjects as part of a complete physiological measurement. The accuracy of the measured mean OEF value over multiple subjects demonstrates its utility as a non-invasive method of estimating baseline CMRO_2_. Mean OEF value across 17 subjects in visual ROI was 0.443 +/- 0.08. These are similar to previously reported OEF values determined using PET. Ishii et al. reported OEF=0.413 +/- 6.1 in the visual cortex; Ibaraki et al. reported 0.39 +/- 0.05 in occipital cortex (Ibaraki et al., 2007; Ishii et al., 1996). The absolute baseline CMRO_2_ calculations were 2.58 +/- 0.53 mM/min (approximately 6.56 ml/100 ml/min) for visual ROI. This is higher than previously reported PET values for baseline CMRO_2_; Ishii et al. reported 4.36 +/- 1.03 ml/100ml/min in visual cortex, while Ibaraki et al. reported 4.3 +/- 0.7 ml/100ml/min in occipital cortex (Ibaraki et al., 2007; Ishii et al., 1996). Part of this difference is most likely due to a higher baseline CBF in our study due to the active baseline. There are multiple components to the active baseline—a visual component, as the subject is fixating on the changing numbers, an attention/memory component, as the subject is focused on the order of the numbers, and a motor component in the form of button presses. As a result, the higher CBF baseline is likely due to the involvement of the task, as compared to a resting baseline with the subject simply lying still. Baseline CBF was determined to be 72.7 +/- 11.8 ml/100ml/min in visual ROI. This is compared to 66.1 +/- 12.9 in visual cortex (Ishii et al., 1996), or 58 +/- 11 ml/100ml/min in occipital cortex (Ibaraki et al., 2007), as reported previously. In addition, we observed a striking difference in the CBF between our female and male subjects, and considering the large effect size equaling 1.5 standard deviations, we feel confident in concluding that female CBF is greater than male CBF overall. While a difference in hematocrit between males and females could affect blood T_2_* and thus produce an artefactual difference in the ASL measurements, the data here were collected with a short echo time that minimized such an effect (< 2%).

Interestingly, the correlation between baseline OEF and baseline CBF was weak (i.e., small effect) and nonsignificant. It is possible that a baseline OEF around 40% is in some sense optimal, and that across subjects, baseline CBF adjusts to produce this OEF for the CMRO_2_ demands of the individual. The distribution of OEF measurement across the group was thus much tighter than CBF measurements, which has been documented previously using other techniques (He and Yablonskiy, 2007; Raichle et al., 2001). Additionally, the variance observed is reflective of the variance in measurement technique combined with true variance across subjects. Because of the uniformity of the subject population, we did not expect large physiological variations in OEF across the subject pool. We thus could assume that the variance observed is dominated by the variance in measurement technique. The variance across subjects in measuring OEF with VSEAN compared to the variance using PET measurements was similar, or even better, depending on the study, as noted above. VSEAN could thus be a comparable technique to utilize that does not require the invasiveness of PET.

At this point, further development of these techniques will allow clinical researchers to assess how regional metabolism is altered in disease states. VSEAN would be a useful tool in using group mean results to determine changes in CMRO_2_ in pathophysiology. Studies of human development, healthy aging, and disease would benefit from being able to assess group differences in baseline CMRO_2_. There are limitations, however, to the types of diseases to which the methods could be applied; underlying changes to blood flow, blood content, and vasculature in some medical conditions could violate certain assumptions of the acquisition techniques. The pathophysiology of each disease process should be considered in the interpretation of these measurements, and more work is needed to understand how to best apply these methods in clinical applications. A subsequent paper will address the utility of individual and group OEF measurements in detecting CMRO_2_ change in a single subject after significant alteration with drug administration.

Potential limitations of VSEAN include (1) low signal-to-noise (SNR) in certain voxels and (2) measurements that rely on a calibration curve developed using *ex vivo* bovine blood. With regard to (1) low SNR, multi-echo spin echo measurements allow for higher temporal resolution and efficiency, but the accuracy of the T_2_ measurement may be affected by flow artifacts through the multi-echo gradients echo-refocusing effects, and wash-out effects (Guo and Wong, 2012). To combat this, OEF is not calculated on a voxel-wise basis, and instead a T_2_ fit is calculated for the average signals for each echo across a pre-defined mask. To address (2) the use of an *ex vivo* bovine blood calibration curve, while a T_2_-Y calibration curve has been published using human blood samples by Lu and colleagues (Lu et al., 2012), this curve was not used due to a different T_2_-preparation module (hard composite pulses vs. BIR-4 based), a different way of modulating eTEs (increasing the number of pulse modules vs. increasing the gap in the module), and its limited range. The authors noted that the relevant oxygenation range for venous blood, 0.5-0.75, was not tested (Lu et al., 2012); since VSEAN was developed specifically to measure blood oxygenation at those lower levels, the calibration curve using human blood was forgone for the Zhao curve that has a range of 0.39-1.00 for Hct=0.44. We would, however, still expect an effect of hematocrit on T_2_ values as shown by previous investigators (Golay et al., 2001; Lu et al., 2012), and thus the estimate would most benefit from a Y-Hct dependent calibration curve, were it available, to control for the effect of hematocrit.

### 4.2 ΔCMRO_2_ from FLAIR-GESSE R_2_′-M calibration

Like VSEAN, previous studies have not evaluated FLAIR-GESSE to quantify CMRO_2_ in a large sample of subjects. Here, we assumed the ideal relationship *M′* ≅ TE • R_2_*′* to calculate *M′* values that we then used to calculate *δ*CMRO_2_ to a stimulus. An interesting aspect of our findings is the apparent difference in the relationship of baseline CBF to CMRO_2_ versus in response to visual activation (Figs. 5A and 5B). We did not find a significant relationship between CBF and CMRO_2_ in the baseline condition (Fig. 5A); however, the medium effect size of the relationship (Cohen’s *d* = 0.97) suggests it would achieve significance with a slightly larger sample. In contrast, there was no ambiguity in the strength of the positive relationship between CBF and CMRO_2_ in response to visual activation (Fig. 5B), which was a very large and significant effect (*d* = 5.23). Overall, our results provide clear evidence for a physiologic difference in the coupling of flow and metabolism at baseline versus with activity. At baseline, the coupling is modest, whereas there is tight coupling during activation.

Empirically, our group estimate of *M′* closely resembled the *M* calculated from a hypercapnia experiment, with *α*=0.2 and *β*=1.3. However, both *M* and *M′* depend on assumptions employed in the modeling that complicate direct physiological interpretation of these parameters (Simon et al., 2016). The value of *M* estimated from a hypercapnia experiment depends on the version of the BOLD model used for the analysis, specifically the assumption of values for *α* and *β*, and also the assumption that there is no CMRO_2_ change with hypercapnia. For the newer method, the estimate of R_2_*′* may depend on the acquisition technique and analysis, because the signal decay described by R_2_*′* is not a simple mono-exponential decay. Furthermore, theoretical modeling suggests that *M* is expected to be larger than *M′* due to diffusion and other BOLD effects associated with activation that are not captured by baseline R_2_*′*, such as blood volume changes or intravascular signal changes (Blockley et al., 2015). In practice, uncorrected field distortions, particularly near the edges of the brain where field distortions are likely to be the most nonlinear (Dickson et al., 2010), would tend to make larger estimates of R_2_*′* and thus *M′*. Additionally, different acquisitions have been shown to yield different values of R_2_*′*; work by Ni and colleagues showed GESSE returned lower GM R_2_*′* measurements than asymmetric spin echo (ASE) and lower WM R_2_*′* than both ASE and gradient-echo sampling of free induction decay and echo (GESFIDE) (Ni et al., 2015). A scaling of *M′* may be necessary depending on the acquisition method used. In short, whether one is using *M′* or *M* to calibrate a BOLD/ASL experiment and measure CMRO_2_, one needs to be mindful of the potential for systematic effects on accuracy due to the modeling assumptions and acquisition methods. The current results support the feasibility of doing a calibrated experiment without hypercapnia, but further work is needed to clarify the precise physical and physiologic effects underlying *M* and R_2_*′* and their relationship.

As a test of the statistical properties of the standard hypercapnia method and the newer R_2_*′* method, we compared the distributions of *δ*CMRO_2_ values across the population estimated in two ways: first, using the individually measured values of *M* or *M′*, and second, with the assumption that each subject’s value of *M* or *M′* was the same and equal to the group mean value. The rationale for this comparison is that in general we expect that true variation of the physiological effects modeled by *M* would lead to increased variance of the *δ*CMRO_2_ values estimated across a population, if not taken into account, and use of individual measurements of *M* should then reduce the variance of *δ*CMRO_2_ compared to using a constant value of *M* for all subjects. However, if there is significant random error in the measurement of *M*, and the true physiological range of the physiological parameters across the population is narrow, using individual *M* values could increase the range of estimated *δ*CMRO_2_ across the population due to that random error. A conservative assumption is that the physiological factors that determine *M* and *δ*CMRO_2_ to the stimulus are relatively uniform across our sample of young, healthy control subjects, and thus the observed variance of the estimates of *δ*CMRO_2_ across the population is likely a reflection of measurement error of *M*. The mean and standard deviation of the estimates of *δ*CMRO_2_ calculated using individual *M′* values closely matched changes calculated using the group mean *M′*, consistent with this assumption and only a small role of random error of individual *M′* estimates. However, for the hypercapnia data the same comparison showed that the distribution *δ*CMRO_2_ estimates from individual *M* values had a mean reduced by more than a factor of two, which may be due to the interaction of noisy signals with the nonlinearity of the calculation of *M* using Eq [1]. In addition, the variance increased by about a factor of two compared with the *δ*CMRO_2_ distribution estimated from the group mean *M*, consistent with high variability of the individual *M* measurements. It may be that the higher variance of *δ*CMRO_2_ estimates with individual *M* values is due to our method of CO_2_ administration, and it is possible that real-time end-tidal forcing could yield more consistent results on an individual basis. However, dedicated equipment to enable end-tidal forcing further restricts the application of quantitative physiology methods to dedicated research laboratories, supporting the push to work toward replacing gas-based techniques with gas-free based techniques.

Finally, it is important to note that R_2_*′* measurements, like the OEF measurements with VSEAN, may be most useful in fMRI studies of development, aging and disease in which different groups are compared. For example, a different mean R_2_*′* for children and adults in the brain region associated with performing a particular task could indicate that a difference in the BOLD response between the two groups should be interpreted with caution; the BOLD response difference may reflect a difference in baseline conditions rather than a difference in neural activity. Even if there is uncertainty about the exact relationship between *M* and *M′*, as discussed above, simply normalizing the BOLD response by the mean R_2_*′* for the group would remove the baseline dependence, and any remaining difference between the two groups could be interpreted as due to differences in the CMRO_2_ and CBF responses to the stimuli. If the CBF response also is measured, this will enable interpretation of whether the CMRO_2_ response is different in the two groups based on the measured R_2_*′*.

### 4.3 Voxel-wise versus basic ROI-wise analyses

Our primary analysis of the data focused on a functionally localized ROI in visual cortex. In this approach, all the raw data for voxels within the ROI were first pooled before calculating the physiological variables. That is, fractional CBF and BOLD responses were calculated after pooling the raw voxel data, and OEF and R_2_*′* were calculated after pooling the raw data for voxels within the ROI. This approach was used to provide a basic proof of concept of estimating absolute physiological parameters for both baseline and activation states. In principle, this approach could be extended to voxel-wise analysis, although this will require further developmental work. There are two central problems: 1) the VSEAN measurement suffers from poor estimation of R_2_ when the signal to noise ratio is low; and 2) the R_2_*′* measurement is likely to be positively biased in regions where large-scale field correction is incomplete. For this reason, not every voxel in the visually activated ROI and GM masks may contain accurate information on OEF and R_2_*′*. As a first step in beginning to extend these measurements to individual voxels, we chose limits on the quality of fit of R_2_ in the VSEAN experiment and on the range of R_2_*′* as markers of data of sufficient quality to make an estimate of the associated parameter for that voxel. We then tested how many of the voxels chosen to be in the visual cortex ROI based on the activation data also had good quality estimates of OEF and R_2_*′*. While a relatively large fraction of ROI voxels had a full set of measurements, there is clearly room for more work to improve the robustness of these measurements. Importantly, when we performed individual voxel analyses on just these voxels and then averaged over the calculated values, the basic physiological parameter estimates were similar to the ROI-approach of first pooling the data: the mean or median derived from voxel-wise analysis were not statistically different from the mean OEF and R_2_*′* derived using the basic visual ROI mask. In the future, simply using a visual ROI determined from the functional localizer and GM mask from the ASL flow data may be enough to make representative estimates of OEF and *M*.

### 4.4 Physiological results

The primary goal of this work was to begin to evaluate methods for making quantitative physiological measurements without requiring the subject to breathe special gas mixtures. To that end, a homogeneous group of young adult subjects was studied, and we did not expect there to be large physiological variation across subjects. We found no statistically significant correlations between baseline OEF and baseline CBF, nor between baseline OEF and *M* or *M′*. For the latter, we expect *M* to scale with total deoxyhemoglobin in the baseline state, and total deoxyhemoglobin should scale with the product of OEF and baseline venous blood volume. Part of the variability of *M* may be due to variability of blood volume. Further work is needed to test these methods in a wider range of subjects where more physiological variation could be expected. Looking at sex differences in the measured physiological parameters, we found baseline CBF to be significantly higher in females compared to males by about 22%. One of the potential advantages of a fully quantitative assessment of CBF and CMRO_2_ in both the baseline and activation states is that we can examine absolute changes to a stimulus in addition to the more commonly measured fractional changes. In comparing absolute baseline values of CBF and CMRO_2_ and absolute changes in these quantities to the stimulus, we found a different slope (Figure 5A and 5B). This suggests that different physiological mechanisms may serve to balance baseline CBF/CMRO_2_ coupling and activation CBF/CMRO_2_ coupling, although more work is needed to define these mechanisms.

## 5. Conclusions

The results from this work demonstrate the potential for making absolute CMRO_2_ measurements without requiring the subject to breathe special gases. However, work is still needed to refine these methods and establish their validity. Each method has its own challenges: for VSEAN, low SNR of the isolated blood signal for some voxels, and for FLAIR-GESSE, excessive field distortion near edges of the brain. Nevertheless, these results support the feasibility of painting a more complete picture of brain physiology, in absolute terms, during both baseline and activation states, with the potential of providing a deeper probe of neural activity in the human brain.

## Acknowledgements

This work was supported by the National Institutes of Health grants NS036722, MH111359, MH112969, and MH113295.

The authors acknowledge and thank Kenny Jackson, Abel Martinez, Jessica Ho, and David Shin for their contributions to this project.

## Abbreviations

BOLD: Blood Oxygenation Level Dependent
ASL: Arterial spin labeling
PICORE: Proximal inversion with control of off-resonance effects
QUIPSS II: Quantitative imaging of perfusion using a single subtraction II
VSEAN: Velocity-Selective Excitation and Arterial Nulling
FLAIR-GESSE: FLuid Attenuated Inversion Recovery Gradient Echo Sampling of Spin Echo

## Data and code availability

The data that support the findings of this study are available from the corresponding author, RBB, upon reasonable request. The data and code sharing adopted by the authors comply with requirements by the National Institutes of Health and by the University of California, San Diego, and comply with institutional ethics approval.

